# LEDGF interacts with the NID of MeCP2 and modulates MeCP2 condensates

**DOI:** 10.1101/2024.04.24.590897

**Authors:** Saskia Lesire, Rodrigo Lata, Yannick Hoogvliets, Kune Herrebosch, Paulien Van De Velde, Anouk Speleers, Frauke Christ, Siska Van Belle, Zeger Debyser

**Affiliations:** KU Leuven, Department of Pharmaceutical and Pharmacological Sciences, Leuven, Flanders, Belgium

**Keywords:** MeCP2, LEDGF, heterochromatin, Rett Syndrome

## Abstract

Methyl-CpG-binding protein 2 (MeCP2) is a ubiquitously expressed nuclear protein that is involved in transcriptional regulation and chromatin remodeling. MeCP2 exists in two isoforms, MeCP2 E1 and MeCP2 E2, which share the same functional domains. Loss-of-function mutations in the MeCP2 gene are the main cause of Rett syndrome (RTT). Previous studies identified a direct interaction between MeCP2 and Lens Epithelium-derived Growth Factor (LEDGF), a transcriptional regulator that also exists in two isoforms, LEDGF/p75 and LEDGF/p52. Here, we further characterized the molecular and functional interaction between MeCP2 and LEDGF. The NID domain in MeCP2 is crucial for the binding to the PWWP-CR1 region of LEDGF. Introduction of R306C, a known RTT mutation in the NID of MeCP2, reduced the interaction with LEDGF. Our data reveal mutual inhibition of MeCP2 and LEDGF multimerization due to overlapping binding sites. In line with this observation, LEDGF depletion resulted in enlarged MeCP2 and heterochromatin condensates in NIH3T3 cells. Unraveling the molecular interaction and functional impact of the MeCP2-LEDGF interaction will increase our understanding of RTT pathogenesis.

## Introduction

The methyl-CpG-binding protein 2 (MeCP2) is a ubiquitously expressed nuclear protein that is involved in transcriptional regulation and chromatin remodeling^1^. Differential splicing results in two isoforms, MeCP2 E1 and MeCP2 E2 that differ in function and expression profile (Figure 1A)^2,3^. MeCP2 E1 is encoded by exon 1, 3 and 4, while MeCP2 E2 is encoded by exon 2, 3 and 4, resulting in a 21 amino acid difference at the start of the protein^3^. Both isoforms share the same functional domains, yet they differ in their N-terminal domain (NTD). The NTD is followed by a Methyl-binding Domain (MBD) that binds 5-methyl cytosine (5mC) and 5-hydroxymethyl cytosines (5hmC), an intervening domain (ID) and a transcriptional repression domain (TRD). Within the TRD, at the C-terminus, the NCor Interaction Domain (NID) functions as a recruitment platform for MeCP2’s multiple binding partners^4^. Additionally, MeCP2 contains three AT-hook domains, in the ID, TRD and CTD, that can simultaneously bind DNA independently of its methylation status^1^.

**Figure 1:**
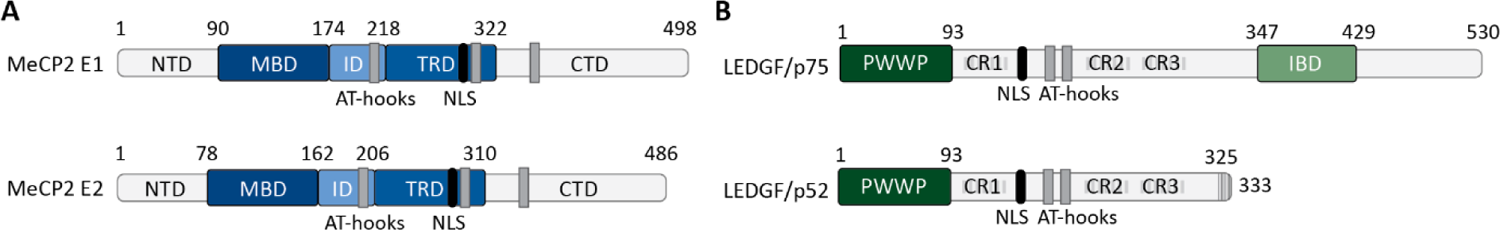
Structure of MeCP2 and LEDGF isoforms. **A.** Structure of MeCP2 E1 and MeCP2 E2 ^5^. NTD: N-terminal Domain, MBD: Methyl-binding Domain, ID: Intervening Domain; TRD: Transcriptional Repression Domain, CTD: C-terminal Domain. **B.** Structure of LEDGF/p75 and LEDGF/p52 ^6^. PWWP: Pro-Trp-Trp-Pro domain, IBD: Integrase Binding Domain, CR: Charged Regions, NLS: Nuclear Localization Signal.

While the two isoforms are abundantly expressed in the central nervous system, different expression levels and distributions are found in developing and post-natal mouse brains^3^. MeCP2 E1 is the major protein isoform, with a relatively uniform expression across the brain. MeCP2 E2, on the other hand, displays a later expression onset during development and shows differential enrichment across different brain regions^3^. When looking at the amino acid sequence, the two isoforms only differ in the NTD, which determines the turn-over rates of MeCP2 as well as the ability of the MBD to interact with DNA^2^. The spatial and temporal difference in MeCP2 isoform expression suggest distinct roles for the two isoforms.

Proper functioning of MeCP2 is essential for normal development and function of the nervous system^7^. MeCP2 is an epigenetic reader for DNA methylation marks and binding of MeCP2 to methylated DNA either silences or promotes gene expression, depending on its interaction partners^8,9^. MeCP2 is also involved in the modification of chromatin structure through the binding of methylated histone H3K9 and H3K27 marks^49^. By recruiting chromatin-modifying proteins, also known as ‘writers’, such as histone deacetylases and methyltransferases, binding of MeCP2 to nucleosomes induces condensation of heterochromatin^9,10^.

Loss-of-function mutations in the MeCP2 gene are the main cause of Rett syndrome (RTT), a neurodevelopmental disorder that affects 1 in 10,000 – 15,000 females^11^. MeCP2 duplication syndrome (MDS) is the mirror disease of RTT. MDS is caused by a duplication of the MeCP2 gene, leading to an overexpression of MeCP2^12^. Most MDS patients are boys, affecting 1 in 150,000 births. Currently, there is no cure for RTT or MDS, and treatment consists of symptomatic therapies to slow down the progression of the disease and improve the life quality of the patients.

A study of the interactome of MeCP2 in the mouse brain revealed that MeCP2 directly interacts with Lens Epithelium Derived Growth Factor (LEDGF), another important regulator of gene transcription^16^. This finding confirmed an earlier study where a protein-protein interaction was identified *in vitro* between MeCP2 and LEDGF^17^. The latter study showed an interaction between MeCP2 and the N-terminal region of LEDGF that is shared between the two LEDGF isoforms. LEDGF is a ubiquitously expressed DNA-binding protein that functions as a transcriptional co-activator by interacting with the RNA polymerase II complex^18^. LEDGF has two isoforms, LEDGF/p75 and LEDGF/p52, which are generated through alternative splicing but share the N-terminal part^19^. Both isoforms contain a PWWP domain that allows LEDGF to interact with DNA regions that contain specific histone modifications^20^. In particular, LEDGF interacts with di-and trimethylated H3K36 marks, tethering regulatory complexes to actively transcribed genes and contributing to the definition of chromatin states and regulating gene expression^21,22^. This binding to chromatin is supported by three charged regions (CR1, CR2 and CR3) and two AT-hooks C-terminally of the PWWP domain. The longer isoform LEDGF/p75 contains an additional Integrase Binding Domain (IBD) which serves as an interaction site for cellular binding partners such as MLL1, PogZ, JPO2 and IWS1^23^.

In this study we determined the molecular and functional interaction between MeCP2 and LEDGF. We first characterized the interaction domains in both MeCP2 and LEDGF, and investigated the possible differences between their respective isoforms. Next, we studied the effect of the MeCP2-LEDGF complex on MeCP2 condensates and heterochromatin formation.

## Materials and methods

### Cell culture

HEK293T cells were grown in a humified atmosphere containing 5% CO_2_ at 37 °C. Cells were cultured in Dulbecco’s modified Eagle’s medium (DMEM) with GlutaMAX [Gibco] supplemented with 5% fetal bovine calf serum (FCS; [Gibco]) and 50 μg/mL gentamicin [Gibco]. Cultured cells were routinely checked for mycoplasma using the PlasmoTest Mycoplasma Detection kit [Invivogen].

### Co-immunoprecipitation

For co-immunoprecipitation assays (co-IP) 7 x 10^6^ HEK293T cells were plated in 10 cm^2^ dishes in DMEM with GlutaMAX [Gibco] supplemented with 2% (v/v) FCS [Gibco] and 50 μg/mL gentamicin [Gibco]. After 24 hours cells were transfected with 20 μg of plasmid DNA (pCHMWS_3xFlag-MeCP2_E1 or pCHMWS_3xFlag-MeCP2_E2) using 10 µM branched PEI [Sigma-Aldrich]. Cells were harvested 24 hours after transfection and lysed in RIPA buffer (150 mM Tris-HCl [Sigma-Aldrich], 150 mM NaCl [Sigma-Aldrich], 0.5% (w/v) sodium deoxycholate [Merck Life Science BV], 0.1% (w/v) sodium dodecyl sulphate (SDS; [Acros Organics], 1% (v/v) IGEPAL [Sigma-Aldrich], pH 8) supplemented with protease inhibitor [cOmplete, EDTA-free, Roche]. The samples were incubated on ice for 30 minutes followed by centrifugation at 21,000 *g* for 10 minutes. In case of Flag-tagged IP, the supernatant was incubated with anti-Flag-beads [Sigma-Aldrich; A2220] and 3 U/mL DNase [Merck Life Science BV] overnight on a turning wheel at 4 °C. The beads were washed using TBS (50 mM Tris-HCl [Sigma-Aldrich], 150 mM NaCl [Sigma-Aldrich], pH 7.4) and collected using centrifugation at 6800 *g*. Immunoprecipitated proteins were detected by western blotting. Input samples and precipitated proteins were separated on a 4-15% Tris-glycine gel [Bio-Rad Laboratories] and electroblotted on Amersham^TM^ Protran Nitrocellulose membranes [VWR]. Membranes were blocked in PBS with 0.1% (v/v) Tergitol [Acros Organics] and 5% (w/v) milk. Subsequently membranes were incubated with primary antibodies (1:500 rabbit anti-LEDGF-PWWP [Abcam; ab177159], 1:1000 rabbit anti-MeCP2 [Cell Signaling Technology; 3456S], 1:1000 rabbit anti-GAPDH [Abcam; Ab9485], 1:400 rabbit anti-Flag [Sigma-Aldrich; F7425]). Detection was performed using secondary horseradish peroxidase-conjugated goat anti-rabbit [Agilent] and chemiluminescent substrate Clarity ECL or Clarity Max ECL [Bio-Rad Laboratories]. Imaging was done with the Amersham^TM^ ImageQuant 800 Western blot imaging system [Cytvia]. Quantification was performed in the Image Lab software [Bio-Rad Laboratories] and statistical analysis was performed using GraphPad Prism 10.0 software package.

Alternatively, cells were transfected with 20 μg of pCHMWS_HA-LEDGF/p75 using 10 µM branched PEI [Sigma-Aldrich]. Magnetic anti-HA-beads [MedChemExpress] were used for precipitation of HA-LEDGF/p75. The beads were washed using TBS-T (50 mM Tris-HCl [Sigma-Aldrich], 150 mM NaCl [Sigma-Aldrich], 0.5% Tween-20 [AppliChem], pH 7.4) and collected using a magnetic holder. Precipitated proteins were eluted in SDS buffer (2% (w/v) SDS [Acros Organics], 160 mM Tris-HCl [Sigma-Aldrich], pH 6.8) and analogously detected by western blotting. Primary antibodies used were: 1:1000 rabbit anti-LEDGF-PWWP [Abcam; ab177159], 1:500 rabbit anti-MeCP2 [Cell Signaling Technology; 3456S], 1:1000 rabbit anti-GAPDH [Abcam; Ab9485], rabbit anti-Flag [Sigma-Aldrich; F7425]).

### Protein purification

#### GST-LEDGF/p52 and GST-LEDGF/p52_ΔPWWP-CR1

GST-LEDGF/p52 or GST-LEDGF/p52_ΔPWWP-CR1 were expressed from pGEX_GST-LEDGF/p52 or pGEX_GST-LEDGF/p52_dPWWP-CR1 in *E. coli* BL21pLysS competent bacteria and grown in LB medium medium supplemented with 100 μg/mL ampicillin [Sigma-Aldrich]. Bacterial cultures were grown at 37 °C until an OD_600_ of 0.6 before protein expression was induced by adding 1 mM IPTG [Sigma-Aldrich]. After 4 hours at 37 °C, cultures were harvested by centrifugation for 10 minutes at 2100 *g*. Pellets were resuspended in 20 mL cold STE buffer (10 mM Tris-HCl [Sigma-Aldrich], pH 7.3, 100 mM NaCl [Sigma-Aldrich], 0.1 mM ethylenediaminetetraacetic acid (EDTA; [Sigma-Aldrich]), and stored at −20 °C. The pellet was lysed using lysis buffer (50 mM Tris-HCl [Sigma-Aldrich], pH 7.5, 250 mM NaCl [Sigma-Aldrich], 1 mM dithiotreitol (DTT; [VWR Chemicals])) supplemented with protease inhibitor [cOmplete, EDTA-free, Roche] and lysed further by sonication [SFX250 Sonifier, Branson]. After sonication, 0.1 μg/mL DNase [Roche] was added and the lysate was incubated for 20 minutes on ice. The lysate was cleared by a 30-minute centrifugation at 27,000 *g*. GST-tagged proteins were purified by affinity chromatography on Glutathione Sepharose-4 Fast Flow [GE Healthcare]. The resin was equilibrated with wash buffer (50 mM Tris-HCl [Sigma-Aldrich], pH 7.5, 250 mM NaCl [Sigma-Aldrich], 1 mM DTT [VWR Chemicals]) and bound proteins were eluted in wash buffer supplemented with 20 mM glutathione. The fractions were analyzed by SDS-PAGE and a Coomassie stain [Coomassie Brilliant Blue G250, Merck Life Science BV]. Peak fractions were pooled and dialyzed against 50 mM Tris-HCl, pH 7.5, 250 mM NaCl [Sigma-Aldrich] and 10% (v/v) glycerol [VWR chemicals].

#### Flag-LEDGF/p52 and Flag-LEDGF/p75

Flag-LEDGF/p52 and Flag-LEDGF/p75 were expressed from pCPnat_3xFlag-LEDGF/p52 or pCPnat_3xFlag-LEDGF/p75 in *E. coli* BL21 Star (DE3) competent bacteria and grown in LB medium supplemented with 100 μg/mL ampicillin [Sigma-Aldrich]. Bacterial cultures were grown at 37°C until an OD_600_ of 0.6 before protein expression was induced by adding 0.5 mM IPTG [Sigma-Aldrich]. After 4 hours at 29 °C, cultures were harvested by centrifugation for 10 minutes at 2100 *g*. Pellets were resuspended in 20 mL cold STE buffer (10 mM Tris-HCl [Sigma-Aldrich], pH 7.3, 100 mM NaCl [Sigma-Aldrich], 0.1 mM EDTA [Sigma-Aldrich]), and stored at −20 °C. The pellet was lysed using lysis buffer (50 mM Tris-HCl [Sigma-Aldrich], pH 7.5, 250 mM NaCl [Sigma-Aldrich], 1 mM DTT [VWR Chemicals]) supplemented with protease inhibitor [cOmplete, EDTA-free, Roche] and lysed further by sonication [SFX250 Sonifier, Branson]. After sonication, 0.1 μg/mL DNase [Roche] was added and the lysate was incubated for 20 minutes on ice. The lysate was cleared by a 30-minute centrifugation at 27,000 *g*. The supernatant was filtered through a 0.22 μm Millex-GS Syringe Filter Unit [Merck Life Science BV] before purification over a 5 mL HiTrap Heparin HP Column [Cytiva], equilibrated in 150 mM NaCl [Sigma-Aldrich], 30 mM Tris-HCl [Sigma-Aldrich] pH 7.0 and 1 mM DTT [VWR Chemicals]. The protein was eluted by increasing the salt concentration from 150 mM up to 2 M NaCl [Sigma-Aldrich] using the AKTA purifier [GE Healthcare] and Unicorn v5 software. Peak fractions were analyzed by SDS-PAGE and a Coomassie stain [Coomassie Brilliant Blue G250, Merck Life Science BV]. Fractions containing Flag-LEDGF were pooled and loaded on a superposeTM 6 10/300 GL size exclusion column [GE Healthcare] to purify further. The size exclusion column was equilibrated in 150 mM NaCl [Sigma-Aldrich], 30 mM Tris-HCl [Sigma-Aldrich], pH 7.4 and 1 mM DTT [VWR Chemicals]. Peak fractions were again analyzed on SDS-PAGE followed by a Coomassie stain [Coomassie Brilliant Blue G250, Merck Life Science BV]. The fractions containing Flag-LEDGF were supplemented with 10% (v/v) glycerol [VWR chemicals] and stored at −80 °C.

#### Flag-MeCP2 E1 and Flag-MeCP2 E2

Full length Flag-MeCP2 E1 and E2 were produced in HEK293T cells. 7 x 10^6^ cells were plated in 10 cm^2^ dishes in DMEM with GlutaMAX [Gibco] supplemented with 2% FCS [Gibco] and 50 μg/mL gentamicin [Gibco]. After 24 hours cells were transfected with 20 μg of plasmid DNA (pCHMWS_3xFlag-MeCP2_E1 or pCHMWS_3xFlag-MeCP2_E2) using 13 µM linear PEI [Polysciences]. Cells were lysed 24h after transfection in lysis buffer (75 mM Tris-HCl [Sigma-Aldrich], 400 mM NaCl [Sigma-Aldrich], 1 mM DTT [VWR Chemicals], pH 8) supplemented with protease inhibitor [cOmplete, EDTA-free, Roche] followed by sonication [SFX250 Sonifier, Branson]. After sonication, 0.1 μg/mL DNase [Roche] was added and the lysate was incubated for 30 minutes on ice. The lysate was cleared by a 30-minute centrifugation at 27,000 *g*. The supernatant was filtered through a 0.22 μm Millex-GS Syringe Filter Unit [Merck Life Science BV] before purification over a 5 mL HiTrap SP HP Column [Cytiva], equilibrated in 30 mM Tris-Base, 200 mM NaCl [Sigma-Aldrich], 1 mM DTT [VWR Chemicals], pH 8. The protein was eluted by increasing the salt concentration from 200 mM up to 2 M NaCl [Sigma-Aldrich] using the AKTA purifier [GE Healthcare] and Unicorn v5 software. Peak fractions were analyzed by SDS-PAGE and a Coomassie stain [Coomassie Brilliant Blue G250, Merck Life Science BV]. The fractions showing the expected molecular weight and purity were pooled and loaded on a superposeTM 6 10/300 GL size exclusion column [GE Healthcare] to purify further. The size exclusion column was equilibrated in 30 mM Tris-Base, 2 M NaCl [Sigma-Aldrich], 1 mM DTT [VWR Chemicals], pH 8. The fractions containing Flag-MeCP2 E1 or Flag-MeCP2 E2 were supplemented with 10% (v/v) glycerol [VWR chemicals] and stored at −80 °C.

#### His_6_-MeCP2 E2 and His_6_-MeCP2 fragments

His_6_-MeCP2 E2 and His_6_-MeCP2 fragments were cloned in pET constructs (pET_His_6_-MeCP2_E2, pET_His_6_-MBD-ID-TRD, pET_His_6_-MBD-ID, pET_His_6_-ID-TRD, pET_His_6_-MBD, pET_His_6_-ID, pET_His_6_-ID-TRD_K210A-KR211A-K215A-K219A, pET_His_6_-ID-TRD_K266A-K267A-R268A-R270A-K271A, pET_His_6_-ID-TRD_K304A-K305A-R306A-K307A-R309A, pET_His_6_-ID-TRD_R306C). Numbering of the amino acid sequences was based on the MeCP2 E2 isoform (Figure 4A). His_6_-MeCP2 E2 and His_6_-MeCP2 fragments were expressed in *E. coli* BL21 competent bacteria. Bacterial cultures were grown at 37 °C until an OD_600_ of 0.8 in LB medium supplemented 0.5% (v/v) glycerol [VWR chemicals] and 100 μg/mL ampicillin [Sigma-Aldrich]. Protein expression was induced by adding 0.25 mM isopropyl β-D-1-thiogalactopyranoside (IPTG; [Sigma-Aldrich]) and bacterial cultures were grown at 18 °C for 20 hours. Cultures were harvested by centrifugation for 10 minutes at 2100 *g*. Pellets were resuspended in 20 mL cold STE buffer (100 mM NaCl [Sigma-Aldrich] [Sigma-Aldrich], 10 mM Tris-HCl [Sigma-Aldrich] [Sigma-Aldrich] pH 7.4 and 0.1 mM EDTA [Sigma-Aldrich]). Bacteria were lysed using lysis buffer (25 mM Tris-HCl [Sigma-Aldrich], 1 M NaCl [Sigma-Aldrich], 10 µM EDTA [Sigma-Aldrich], 1 mM DTT [VWR Chemicals]) supplemented with protease inhibitor [cOmplete, EDTA-free, Roche] and lysed further by sonication [SFX250 Sonifier, Branson]. After sonication, 0.1 μg/mL DNase [Roche] was added and the lysate was incubated for 20 minutes on ice. The lysate was cleared by a 30-minute centrifugation at 27,000 *g*. The fusion His_6_-tagged proteins were captured using His-Select Nickel affinity gel [Sigma-Aldrich; P6611] beads and eluted with STE buffer supplemented with 250 mM imidazole. The eluate was fractionated and analyzed by SDS-PAGE with a Coomassie stain [Coomassie Brilliant Blue G250, Merck Life Science BV]. Fractions with high protein content were pooled and concentrated using Amicon filters [Merck Life Science BV; UFC500324]. The concentrated protein was loaded on a Superose 6 10/300 GL size exclusion column [GE Healthcare], attached to AKTA pure [Cytiva]. The column was equilibrated in 30 mM Tris-HCl [Sigma-Aldrich], 400 mM NaCl [Sigma-Aldrich] and 1mM DTT [VWR Chemicals]. The fractions with absorbance peaks were analyzed by SDS-PAGE with a Coomassie stain [Coomassie Brilliant Blue G250, Merck Life Science BV]. Fractions containing the protein of interest were pooled and stored in 10% (v/v) glycerol [VWR chemicals] at −80 °C.

#### Pull-down assay

For pull-down assays 7.0 x 10^6^ HEK293T cells were plated in 10 cm^2^ dishes. After 48 hours the cells were lysed in RIPA buffer (150 mM Tris-HCl [Sigma-Aldrich], 150 mM NaCl [Sigma-Aldrich], 0.5% (w/v) sodium deoxycholate, 0.1% (w/v) SDS [Acros Organics], 1% (v/v) IGEPAL [Sigma-Aldrich], pH 8) supplemented with protease inhibitor [cOmplete, EDTA-free, Roche]. Samples were incubated on ice for 30 minutes followed by centrifugation at 21,000 *g* for 10 minutes. The supernatant was incubated with 8 μg of recombinant Flag-LEDGF/p75 or Flag-LEDGF/p52, anti-Flag-beads [Sigma-Aldrich; A2220] and 3 U/mL DNase [Merck Life Science BV] overnight on a turning wheel at 4 °C. The beads were washed using TBS (50 mM Tris-HCl [Sigma-Aldrich], 150 mM NaCl [Sigma-Aldrich], pH 7.4) and collected using centrifugation at 6800 *g*. Immunoprecipitated proteins were detected on western blot. Input samples and precipitated proteins were separated on a 4-15% Tris-glycine gel [Bio-Rad Laboratories] and electroblotted on Amersham^TM^ Protran Nitrocellulose membranes [VWR]. Membranes were blocked in PBS with 0.1% (v/v) Tergitol [Acros Organics] and 5% (w/v) milk. Subsequently membranes were incubated with primary antibodies (rabbit anti-LEDGF-PWWP [Abcam; ab177159], rabbit anti-MeCP2 [Cell Signaling Technology; 3456S], rabbit anti-GAPDH [Abcam; Ab9485]). Detection was performed using secondary horseradish peroxidase-conjugated goat anti-rabbit [Agilent] and chemiluminescent substrate Clarity ECL or Clarity Max ECL [Bio-Rad Laboratories]. Imaging was done with the Amersham^TM^ ImageQuant 800 Western blot imaging system [Cytvia].

#### AlphaScreen

AlphaScreen assays were performed according to the manufacturer’s protocol [Perkin Elmer]. Briefly, reactions were performed in 25 μl final volume in a 384-well OptiPlate microtiter plates. The reaction buffer contained 25 mM Tris-HCl [Sigma-Aldrich] pH 7.4, 150 mM NaCl [Sigma-Aldrich], 1 mM MgCl_2_ [Sigma-Aldrich], 0,1% Tween-20 [AppliChem], 0.1% (v/v) BSA [Merck Life Science BV] with or without 20 U/mL MNase. In case MNase was added, 2 mM CaCl_2_ [Sigma-Aldrich] was added to the reaction buffer. Varying concentrations of protein were incubated in 15 μl reaction volume at 4 °C for 1 hour. Subsequently, 10 µg/mL of the donor and acceptor beads were added. After incubation for 1 hour in the dark at room temperature, light emission was measured in the EnVision Xcite Multilabel Reader [Perkin Elmer]. A non-linear regression – sigmoidal curve fit with 1/Y^2^ weighting was fitted to the data^24^.

#### Lentiviral vector production

shRNAs to create MeCP2 or LEDGF KD were generated by annealing sense and antisense oligonucleotide sequences (Table S1). The annealed oligos were cloned into the pGAE_SFFV plasmid backbone using the Esp3I restriction enzyme. For lentiviral vector (LV) production, transfer plasmids carrying the shRNA were cotransfected with a pVSV-G envelope plasmid and a pSIV3+ packaging plasmid as previously described^25,26^. Briefly, LV vectors were produced by triple transfection of HEK293T cells using 10 µM branched PEI [Sigma-Aldrich]. Medium was replaced 24 hours posttransfection and supernatant was collected after 48 and 72 hours by filtration through a 0.45 µm pore-size filter [Merck Life Science BV]. The vectors were concentrated by ultracentrifugation [Amicon Ultra-15 centrifugal filter unit, 50 kDa, Merck Life Science BV].

To rescue MeCP2 depletion, Human MeCP2-eGFP E1 or E2 was cloned into the pCHMWS plasmid backbone for LV production using BamHI and BwiWI restriction enzymes. LV production was performed as described above, using triple transfection of the transfer plasmid, pVSV-G envelope plasmid and p8.91 packaging plasmid.

#### Generation of stable cell lines

First, stable MeCP2 KD NIH3T3 cells were generated by lentiviral (LV) transduction and subsequent selection with 5 μg/mL blasticidin [Invivogen]. MeCP2 KD NIH3T3 cells were subsequently transduced with LV vectors carrying shRNA-resistant human MeCP2-eGFP E1 or E2 and selected with 1 µg/mL puromycin [Invivogen]. These cells were again transduced with LV vectors carrying shRNA’s against LEDGF (Table S1) and selected with 100 µg/mL zeocin [Invivogen]. After transduction cells were continuously kept under selection.

#### Western blot

Western blot was performed to determine protein levels in stably transduced NIH3T3 cell lines. Cells were washed twice with PBS and lysed in RIPA buffer (150 mM Tris-HCl [Sigma-Aldrich], 150 mM NaCl [Sigma-Aldrich], 0.5% (w/v) sodium deoxycholate, 0.1% (w/v) SDS [Acros Organics], 1% (v/v) IGEPAL [Sigma-Aldrich], pH 8) supplemented with protease inhibitor [cOmplete, EDTA-free, Roche]. Protein concentrations of whole cell extracts were determined using the BCA protein assay [ThermoFisher Scientific]. Cell extracts containing 30 μg of total protein were separated on a 4-15% Tris-glycine gel [Bio-Rad Laboratories] and electroblotted on Amersham^TM^ Protran Nitrocellulose membranes [VWR]. Membranes were blocked in PBS with 0.1% (v/v) Tergitol [Acros Organics] and 5% (w/v) milk. Subsequently membranes were incubated with primary antibodies (1:1000 rabbit anti-LEDGF-PWWP [Abcam; ab177159], 1:1000 rabbit anti-GAPDH [Abcam; Ab9485]. Detection was performed using secondary horseradish peroxidase-conjugated goat anti-rabbit [Agilent] and chemiluminescent substrate Clarity ECL or Clarity Max ECL [Bio-Rad Laboratories]. Imaging was done with the Amersham^TM^ ImageQuant 800 Western blot imaging system [Cytvia].

#### Immunocytochemistry

2.5 x 10^4^ NIH3T3 cells were seeded in an 8-well chamber slide [Ibidi] in DMEM with GlutaMAX [Gibco] supplemented with 5% FCS [Gibco] and 50 μg/mL gentamicin [Gibco]. After 24 hours, cells were fixed with 4% (v/v) paraformaldehyde PFA [Sigma-Aldrich] for 20 minutes. Cells were permeabilized using 0.1% (v/v) Triton X-100 [Acros Organics] in PBS followed by 30 minutes incubation in blocking buffer (0.5% (w/v) BSA [Merck Life Science BV] and 0.1% (v/v) Tween-20 [Acros Organics]) at room temperature. Cells were incubated with 1:500 primary rabbit anti-MeCP2 antibody [Cell Signaling Technology] overnight at 4 °C. Cells were incubated for 1 hour at room temperature with the secondary antibody, 1:1000 donkey anti-rabbit Alexa fluor 488, and 5 µg/mL Hoechst [ThermoFisher Scientific]. Images were acquired with the Zeiss LSM 880 confocal microscope at the Cell and Tissue Imaging Core at KU Leuven.

#### High content imaging

96-well black PhenoPlates [Perkin Elmer] were coated using poly-D-lysine (PDL; [Sigma-Aldrich]) and washed twice with PBS before plating 8 x 10^3^ NIH3T3 cells from each cell line in DMEM with GlutaMAX [Gibco] supplemented with 5% FCS [Gibco] and 50 μg/mL gentamicin [Gibco]. 24 hours after plating, cells were synchronized with medium containing 10 µg/mL aphidicolin [ThermoFisher Scientific]. At time intervals, the synchronization medium was replaced with culture medium to measure cells in different stages of the cell cycle. After 30 hours, all cells were fixed with 4% (v/v) PFA [Sigma-Aldrich] and stained with 5 µg/mL Hoechst [ThermoFisher Scientific] for 20 minutes. The plates were imaged using the Operetta CLS High Content Analysis System [Perkin Elmer] at the Bioimaging Core Leuven (VIB-KU Leuven) (channel 1 – wide-field 386-23: Hoechst 333-42, channel 2 – wide-field 488-20: GFP; 20x objective; 4 images per well). The images were analyzed using the Harmony software. First, the nucleus of each cell was detected in the Hoechst channel. The mean area of MeCP2-eGFP speckles and Hoechst speckles inside the nucleus was measured in number of pixels. Results are expressed as mean ± standard deviation (SD). Statistical analysis was done using two-way ANOVA followed by Dunnett’s multiple comparison test vs. WT using the GraphPad Prism 8.0 software package.

## Results

### Interaction between LEDGF and MeCP2

LEDGF and MeCP2 each exist in two distinct isoforms, LEDGF/p75 and LEDGF/p52, and MeCP2 E1 and MeCP2 E2, respectively (Figure 1A, 1B). The direct interaction between MeCP2 and the PWWP-CR1 region of LEDGF has been shown before ^16,17^. We now investigated the interactions of the respective isoforms using co-immunoprecipitation (co-IP). In lysates of HEK293T cells overexpressing Flag-MeCP2 E1 or Flag-MeCP2 E2 both LEDGF isoforms were immunoprecipitated by Flag-MeCP2, confirming their interaction (Figure 2A). No difference was observed between MeCP2 E1 and MeCP2 E2 in precipitating LEDGF. Since both isoforms of LEDGF formed a complex with MeCP2, the domain of LEDGF that is responsible for the interaction with MeCP2 is shared by LEDGF/p75 and LEDGF/p52. Interestingly, the relative abundance of LEDGF/p75 and LEDGF/52 isoforms altered upon precipitation. Whereas LEDGF/p75 is on average 5-fold more abundant in HEK293T cells than LEDGF/p52, in the precipitate this ratio was reduced to a ∼1.5-fold difference (Figure 2B). The quantification suggests a higher affinity of MeCP2 for LEDGF/p52 than for LEDGF/p75. We further validated the MeCP2-LEDGF interaction by performing a pull-down assay showing the interaction between recombinant, *E. coli*-expressed, Flag-LEDGF/p75 or Flag-LEDGF/p52 proteins and endogenous MeCP2 (Figure 2C). Additionally, a reverse co-IP in HEK293T cells overexpressing HA-LEDGF/p75 confirmed the interaction between LEDGF/p75 and endogenous MeCP2 (Figure S1A).

**Figure 2:**
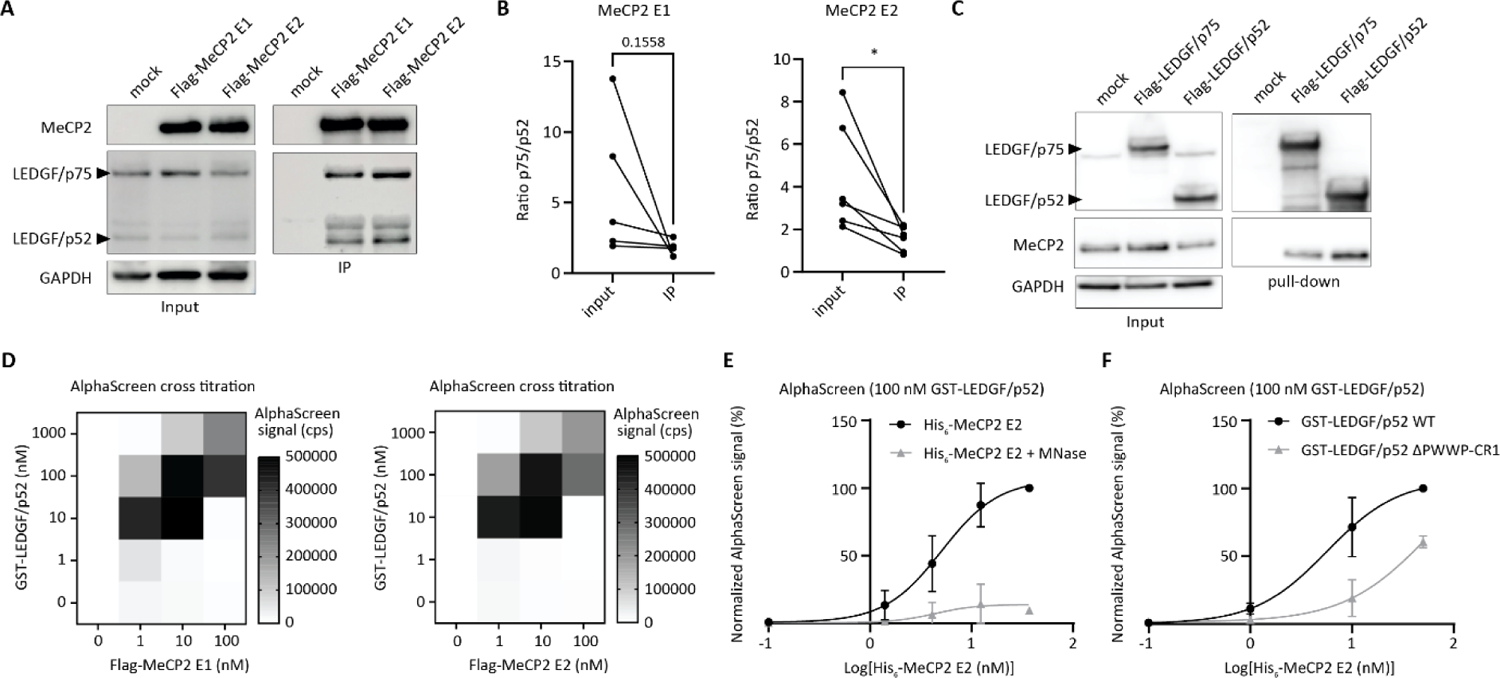
Both isoforms of MeCP2 interact with both isoforms of LEDGF. **A.** Co-IP of MeCP2 E1 and MeCP2 E2. HEK293T cells were transfected with plasmids encoding Flag-MeCP2 E1 or Flag-MeCP2 E2. 24 hours after transfection anti-Flag-beads were used for immunoprecipitation of the cell lysate. Precipitated proteins were analyzed on western blot. MeCP2 was detected with an anti-MeCP2 antibody (1:1000). LEDGF was detected with an anti-LEDGF-PWWP antibody (1:500). GAPDH was detected with an anti-GAPDH-antibody (1:1000). **B.** Quantification of co-IPs. The ratio between LEDGF/p75 and LEDGF/p52 was calculated for the input and IP. Ratios were plotted separately for MeCP2 E1 (n = 5) and MeCP2 E2 (n = 6). Statistical analysis was done using a paired t-test. * p < 0.05. Error bars represent the SD. **C.** Pull-down assay with recombinant Flag-LEDGF/p75 or Flag-LEDGF/p52. 8 µg of recombinant LEDGF/p52 or LEDGF/p52 was added to HEK293T cell lysates. Anti-Flag-beads were used for immunoprecipitation of the cell lysate. Precipitated proteins were analyzed on western blot. LEDGF was detected with an anti-LEDGF-PWWP antibody (1:1000). MeCP2 was detected with an anti-MeCP2 antibody (1:500). GAPDH was detected with an anti-GAPDH-antibody (1:1000). **D.** AlphaScreen cross titration between GST-LEDGF/p52 and Flag-MeCP2 E1 or E2. **E.** AlphaScreen titration between 100 nM GST-LEDGF/p52 and increasing concentrations of His6-MeCP2 E2 with and without MNase treatment. **F.** AlphaScreen titration between 100 nM GST-LEDGF/p52 WT or ΔPWWP-CR1 and His6-MeCP2 E2 without MNase treatment. Error bars represent the SD; n = 3. A non-linear regression – sigmoidal curve fit with 1/Y^2^ weighting was fitted to the AlphaScreen data. See also Figure S1.

To better understand the direct interaction between LEDGF and MeCP2 we performed a series of *in vitro* AlphaScreen experiments. Flag-MeCP2 E1 and Flag-MeCP2 E2 were purified from HEK293T cells. A direct interaction with GST-LEDGF/p52 was demonstrated, with similar apparent affinities for both MeCP2 isoforms (Figure 2D). His_6_-MeCP2 E2 was also expressed in and purified from *E. coli* to determine whether post-translational modifications (PTM) of MeCP2 are required for the binding to LEDGF. Eukaryotic Flag-MeCP2 E2 and bacterial His_6_-MeCP2 E2 showed an identical binding profile to GST-LEDGF/p52, excluding the requirement of PTMs in MeCP2 for its interaction with LEDGF (Figure S1B).

Indirect complex formation between MeCP2 and LEDGF may occur due to the presence of DNA as MeCP2 and LEDGF are both DNA-binding proteins. To remove potential DNA contaminants from protein productions, we performed AlphaScreens in the presence of Micrococcal Nuclease (MNase). Although MNase treatment clearly reduced the observed interaction between His_6_-MeCP2 E2 and GST-LEDGF/p52, a residual DNA-independent interaction was evidenced (Figure 2E).

To determine the interaction domain of LEDGF, a recombinant GST-LEDGF/p52 lacking the PWWP-CR1 domain (amino acids 1-143) was expressed, indicated as the interface by Leoh *et al.*^17^. In line with previously obtained results, the interaction with His_6_-MeCP2 significantly decreased upon deletion of the PWWP-CR1, indicating the importance of this region for the direct interaction with MeCP2 (Figure 2F). This result was confirmed with Flag-MeCP2 E2 (Figure S1C). GST-LEDGF/p52 ΔPWWP-CR1, lacking amino acids 1-143, bound less to Flag-MeCP2 in comparison with WT GST-LEDGF/p52 (Figure S1C). Residual binding of the ΔPWWP-CR1 mutant to MeCP2 may result from indirect complex formation with DNA.

### The NID of MeCP2 interacts with LEDGF

MeCP2 consists of five domains: the N-terminal domain (NTD), the Methyl-binding Domain (MBD), the Intervening Domain (ID), the Transcriptional Repression Domain (TRD) and the C-terminal domain (CTD). To gain a better understanding of the MeCP2-LEDGF interface, we determined which domains of MeCP2 interact with LEDGF by using MeCP2 deletion mutants in a series of co-IPs (Figure 3A). Results showed that the amount of precipitated LEDGF/p75 and LEDGF/p52 was reduced upon deletion of the ID or TRD of Flag-MeCP2, irrespective of the MeCP2 isoform (Figure 3B-C). However, despite the observed reduction in precipitated LEDGF, deletion of the individual ID or TRD did not completely disrupt the interaction. An additional MeCP2 deletion construct was made, removing both the ID and TRD. Co-IPs showed that Flag-MeCP2 ΔID-TRD no longer binds to endogenous LEDGF (Figure 3D-E).

**Figure 3:**
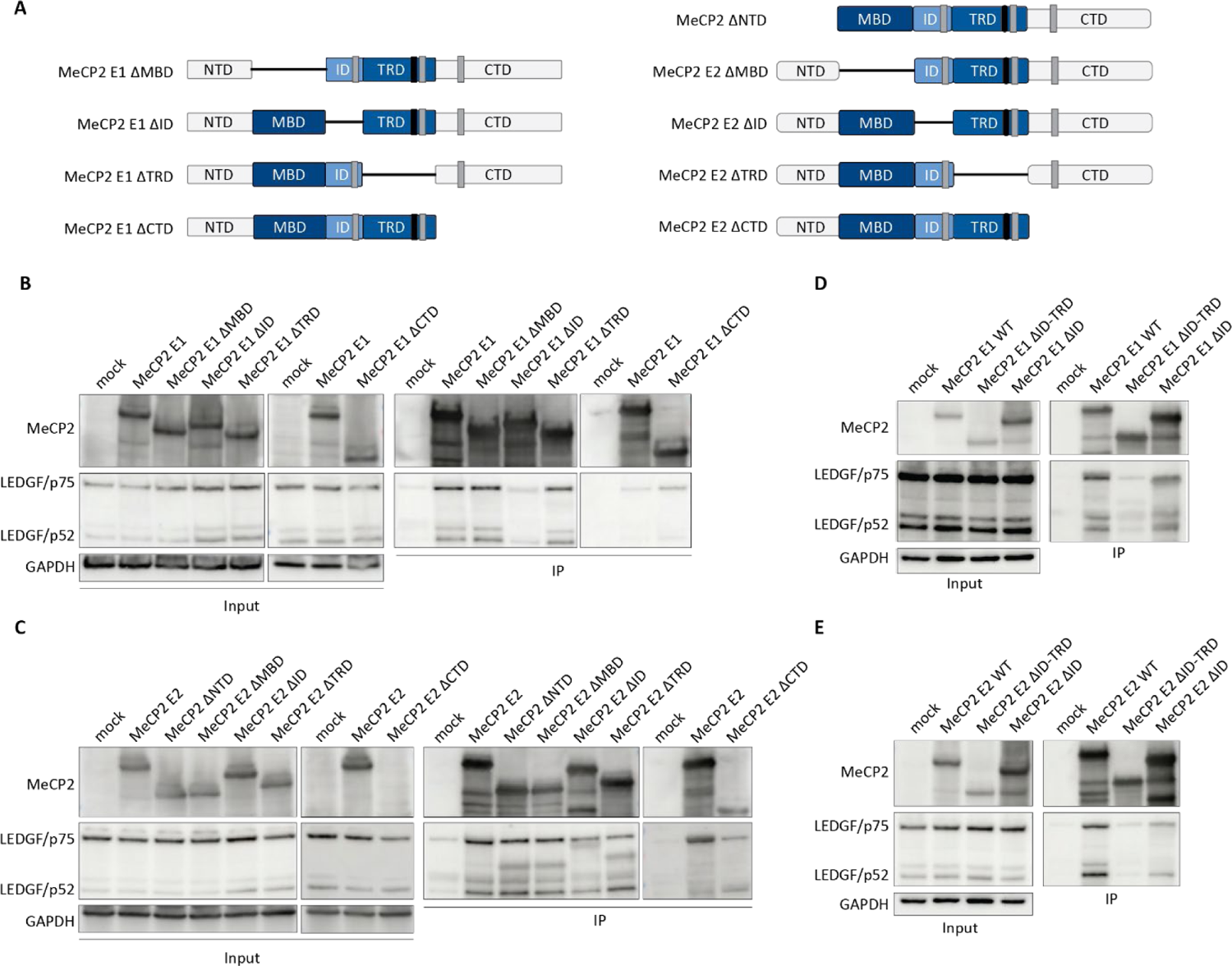
The ID-TRD domain of MeCP2 interacts with LEDGF in HEK293T cells. **A.** Structure of domain deletion constructs of MeCP2 E1 and MeCP2 E2. NTD: N-terminal Domain, MBD: Methyl-binding Domain, ID: Intervening Domain; TRD: Transcriptional Repression Domain, CTD: C-terminal Domain. **B.** Co-IP of MeCP2 E1 deletion constructs. HEK293T cells were transfected with domain deletion plasmids of Flag-MeCP2 E1. 24 hours after transfection anti-Flag-beads were used for immunoprecipitation of the cell lysate. Precipitated proteins were analyzed on western blot. MeCP2 was detected with an anti-MeCP2 antibody (1:1000). LEDGF was detected with an anti-LEDGF-PWWP antibody (1:500). MeCP2-ΔCTD was detected with an anti-Flag-antibody (1:400). GAPDH was detected with an anti-GAPDH-antibody (1:1000). **C.** Co-IP of MeCP2 E2 deletion constructs. HEK293T cells were transfected with domain deletion plasmids of Flag-MeCP2 E2. **D.** Co-IP of MeCP2 E1ΔID-TRD. HEK293T cells were transfected an ID-TRD domain deletion plasmid of Flag-MeCP2 E1. **E.** Co-IP of MeCP2 E2 ΔID-TRD. HEK293T cells were transfected an ID-TRD domain deletion plasmid of Flag-MeCP2 E2. All Co-IPs were performed as described for panel B. Representative western blots are shown (n = 2).

Further *in vitro* AlphaScreens were performed to validate the interaction domain of MeCP2. Recombinant His_6_-MeCP2 fragments were made to study the minimal domain of MeCP2 required for the binding to LEDGF. A His_6_-MBD-ID-TRD construct was made containing only the functional domains of MeCP2 that are shared between the two isoforms (Figure 4A). Removing the NTD and CTD did not change the binding of His_6_-MeCP2 to GST-LEDGF/p52 (Figure S1D). Additionally, we produced fragments corresponding to an individual MeCP2 domain (His_6_-MBD, His_6_-ID) or two consecutive domains (His_6_-MBD-ID, His_6_-ID-TRD) and compared them to the His_6_-MBD-ID-TRD construct. All fragments showed binding to GST-LEDGF/p52 in the presence of DNA (Figure 4B), suggesting DNA-dependence of the observed MeCP2-LEDGF interaction. Therefore, we tested the interaction of the MeCP2 fragments with LEDGF after MNase treatment. When comparing the distinct His_6_-MeCP2 fragments, we observed that the ID-TRD fragment was the only fragment that still binds LEDGF after treatment with MNase (Figure 4B-H). In fact, the DNA-bridging by the ID-TRD domain for binding to GST-LEDGF/p52 was minimal (Figure 4H). This result suggests that the ID-TRD of MeCP2 is responsible for directly binding LEDGF. We reasoned that residual binding of the ΔPWWP-CR1 mutant to His_6_-MeCP2 E2 (Figure 2F) resulted from indirect complex formation with DNA. Therefore, we tested His_6_-ID-TRD against GST-LEDGF/p52 WT or ΔPWWP-CR1 in the presence of MNase. Deletion of the PWWP-CR1 of GST-LEDGF/p52 significantly decreased the binding to His_6_-ID-TRD (Figure 4I).

**Figure 4:**
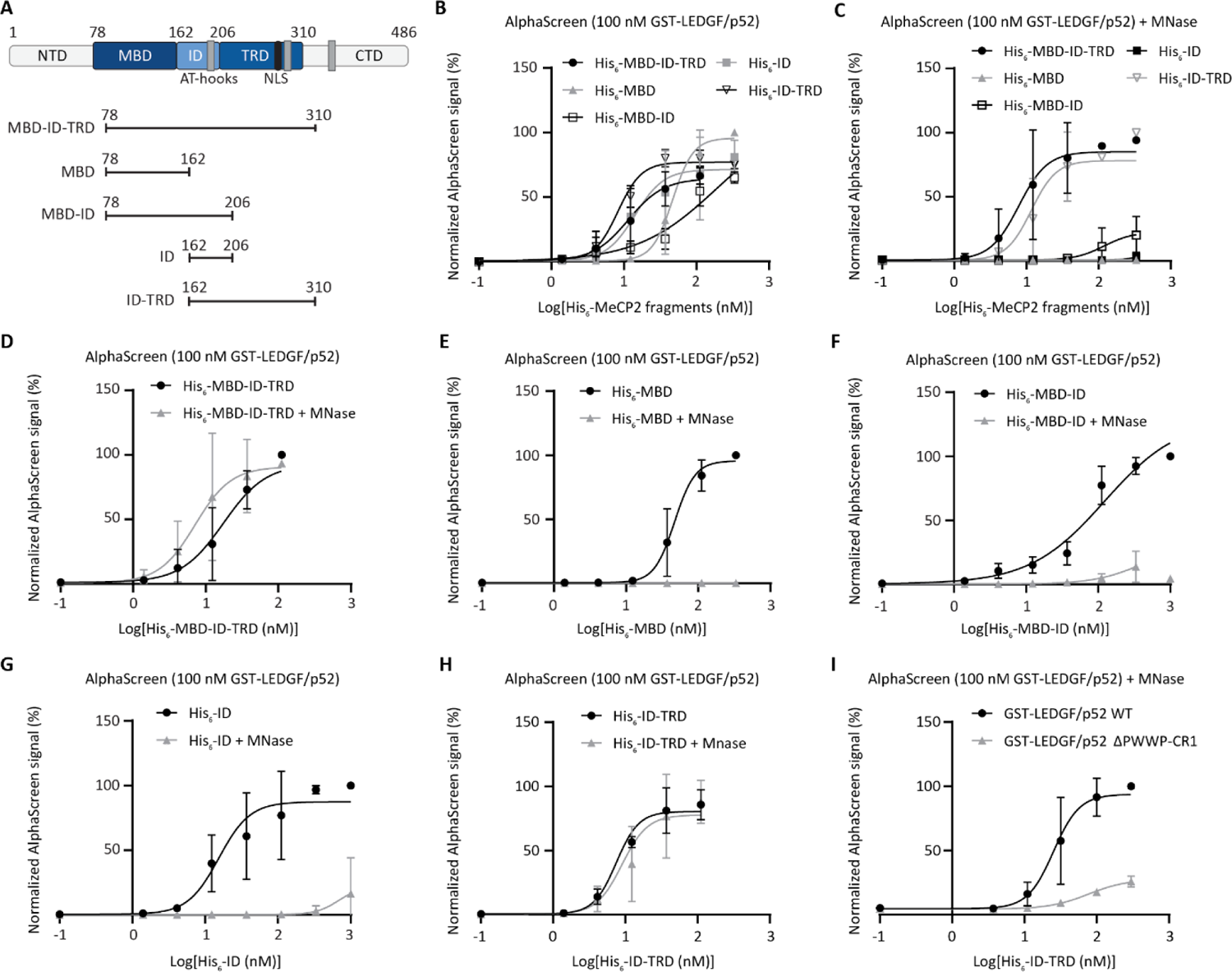
The C-terminal region of the TRD of MeCP2 interacts with the PWWP-CR1 domain of LEDGF. **A**. Schematic representation of recombinant His6-MeCP2 fragments. Numbering is based on the MeCP2 E2 isoform. NTD: N-terminal Domain, MBD: Methyl-binding Domain, ID: Intervening Domain; TRD: Transcriptional Repression Domain, CTD: C-terminal Domain. Following AlphaScreen titrations were performed: **B**. 100 nM GST-LEDGF/p52 and increasing concentrations of His6-MeCP2 fragments without MNase treatment.**C.** 100 nM GST-LEDGF/p52 and increasing concentrations of His6-MeCP2 fragments with MNase treatment**. D.** 100 nM GST-LEDGF/p52 and increasing concentrations of His6-MBD-ID-TRD with and without MNase treatment. **E.** 100 nM GST-LEDGF/p52 and increasing concentrations of His6-MBD with and without MNase treatment. **F.** 100 nM GST-LEDGF/p52 and increasing concentrations of His6-MBD-ID with and without MNase treatment. **G.** 100 nM GST-LEDGF/p52 and increasing concentrations of His6-ID with and without MNase treatment. **H.** 100 nM GST-LEDGF/p52 and increasing concentrations of His6-ID-TRD with and without MNase treatment. **I.** His6-ID-TRD and 100 nM GST-LEDGF/p52 WT or ΔPWWP-CR1 with MNase treatment. Error bars represent the SD; n = 3. A non-linear regression – sigmoidal curve fit with 1/Y^2^ weighting was fitted to the data. See also Figure S1.

To determine which amino acids within MeCP2 are important for binding LEDGF, three sets of alanine mutants were made in three positively charged regions in the TRD (Figure 5A). Mutant one contained four point mutations (K210A, KR211A, K215A, K219A; **mut1**), mutant two contained five point mutations (K266A, K267A, R268A, R270A, K271A; **mut2**), and mutant three contained five point mutations in the NCoR interaction domain (NID) of the TRD (K304A, K305A, R306A, K307A, R309A; **mut3**). Additionally, the R306C mutation was made as it is the most common clinical RTT mutation in the NID of MeCP2 (Figure 5A-B)^27^. We observed a clear reduction in the binding of His_6_-ID-TRD mut2 and mut3 to GST-LEDGF/p52 in the presence of MNase (Figure 5C). The reduction in binding to GST-LEDGF/p52 was less pronounced for His_6_-ID-TRD mut1 that was mutated in the N-terminal region of the TRD (Figure 5C). A combination of mut2 and mut3 in the His_6_-ID-TRD construct resulted in a clear decrease in binding to GST-LEDGF/p52 compared to ID-TRD WT (Figure 5D). In the presence of MNase, the clinical single amino acid R306C RTT mutant bound less to GST-LEDGF/p52 (Figure 5E). In the presence of DNA, all mutants were still able to bind GST-LEDGF/p52 (Figure S1E). Taken together, our results indicate that the NID of MeCP2 is the interface for the interaction with the PWWP-CR1 region of LEDGF.

**Figure 5:**
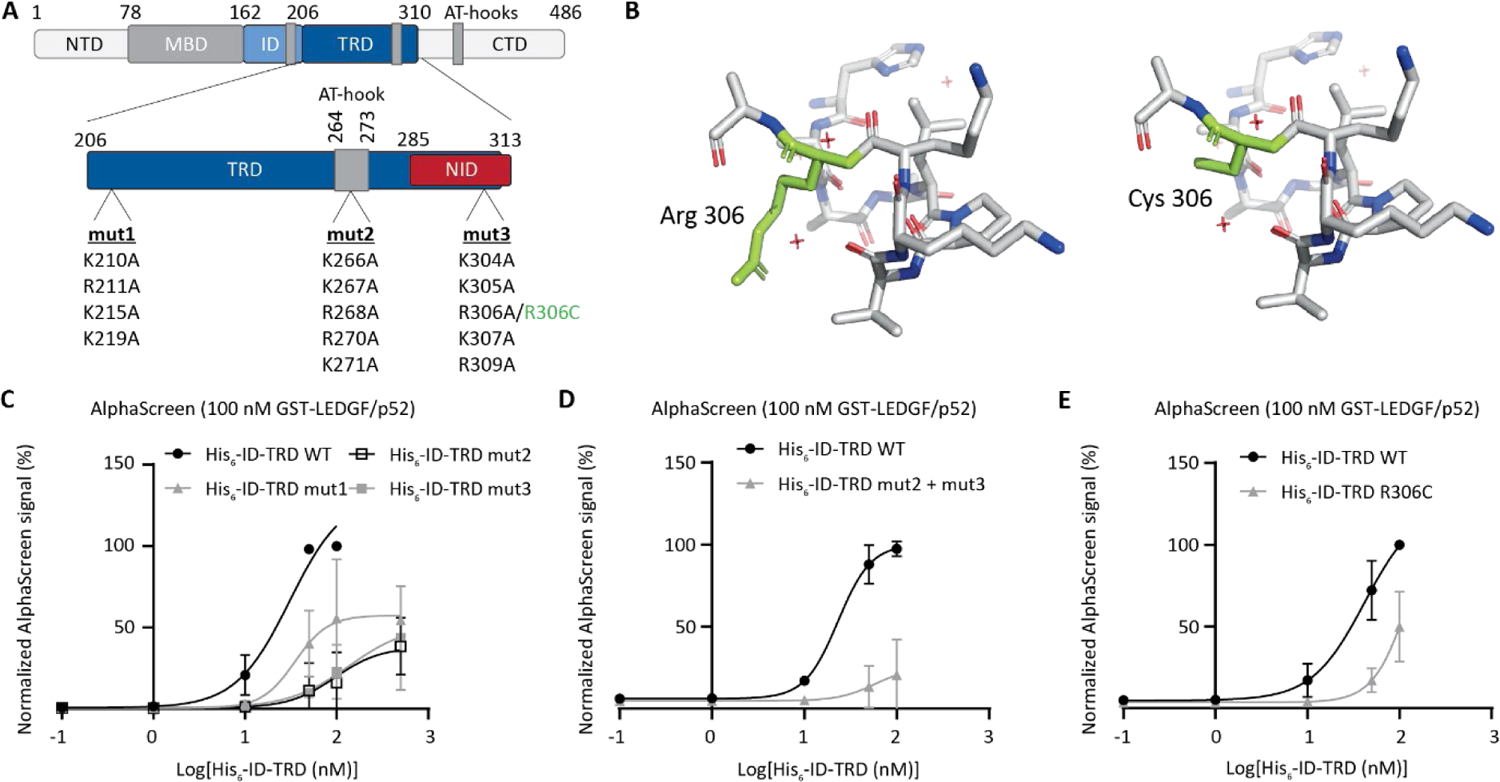
Mutations in the NID of MeCP2 disrupt the interaction with LEDGF. **A.** Structure of MeCP2 with indicated mutations in the TRD. NTD: N-terminal Domain, MBD: Methyl-binding Domain, ID: Intervening Domain; TRD: Transcriptional Repression Domain, CTD: C-terminal Domain. NID: NCoR interaction domain. Mutants were made in the His6-ID-TRD construct. Numbering is based on the MeCP2 E2 isoform. **B.** Structure of the NID of MeCP2 indicating the R306 WT or C306 RTT mutation (PDB ID: 5NAF)^4^. Following AlphaScreen titrations were performed: **C.** 100 nM GST-LEDGF/p52 and increasing concentrations of His6-MeCP2 ID-TRD WT or mutant 1, 2 or 3 with MNase treatment. **D.** 100 nM GST-LEDGF/p52 and increasing concentrations of His6-MeCP2 ID-TRD WT or mutant 2 + 3 with MNase treatment. **E.** 100 nM GST-LEDGF/p52 and increasing concentrations of His6-MeCP2 ID-TRD WT or R306C RTT mutant with MNase treatment. Error bars represent the SD; n = 3. A non-linear regression – sigmoidal curve fit with 1/Y^2^ weighting was fitted to the AlphaScreen data. See also Figure S1.

### MeCP2 homo-oligomerization is disrupted by LEDGF

MeCP2 is known to form electrostatic self-interactions which are essential for heterochromatin formation^28^. Considering that MeCP2 forms multimers and that MeCP2 interacts with LEDGF, we studied the effect of LEDGF on MeCP2 multimerization. We confirmed that Flag-MeCP2 self-associates using *in vitro* AlphaScreens and that this interaction can be disrupted by outcompetition with His_6_-ID-TRD (Figure 6A-B, Figure S1F)^29^. No significant difference in outcompetition with His_6_-ID-TRD WT was seen for the His_6_-ID-TRD R306C RTT mutant (Figure 6C). However, Flag-MeCP2 E2 multimers were disrupted by increasing concentrations of GST-LEDGF/p52 (Figure 6D).

**Figure 6:**
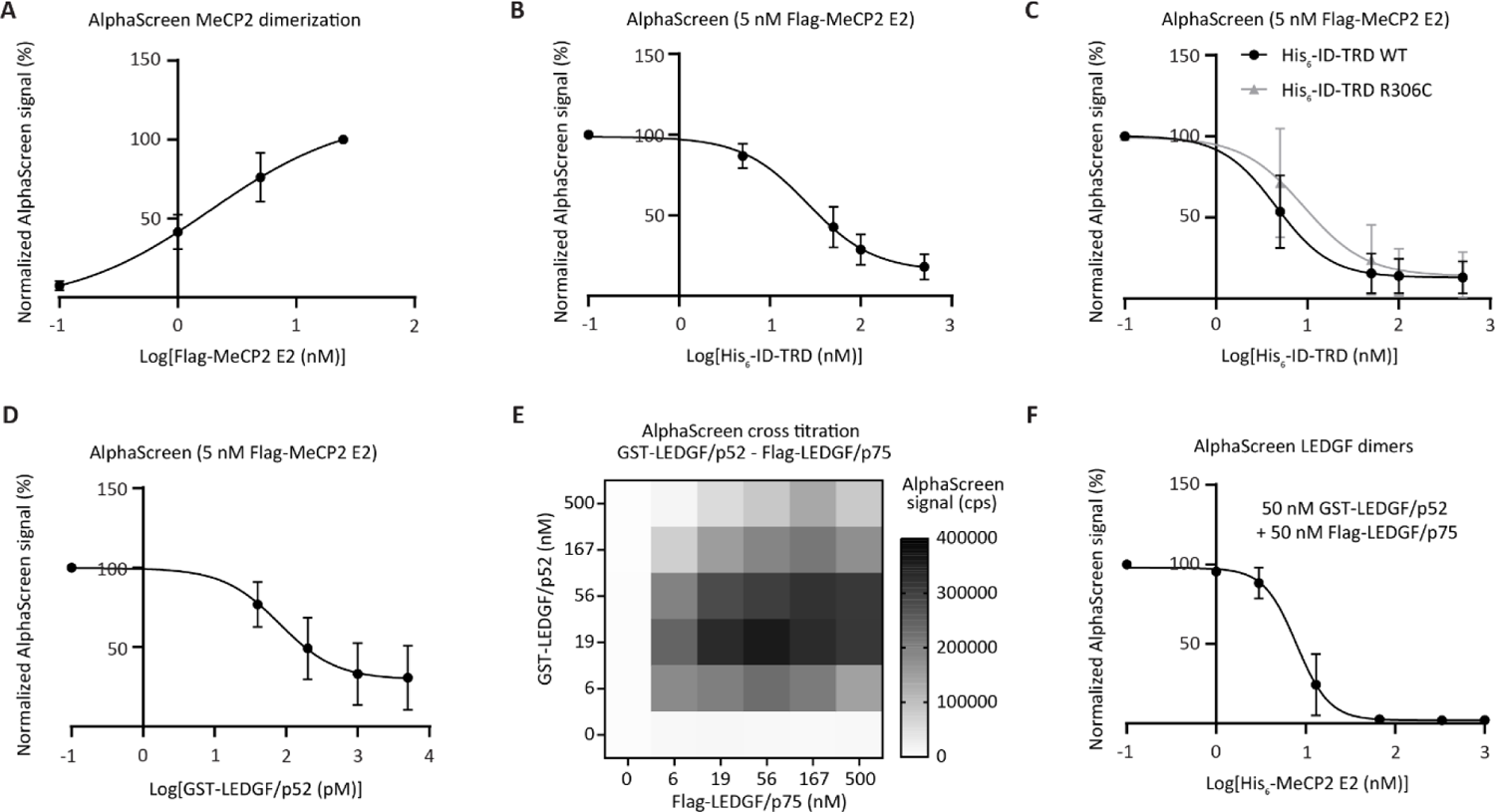
Reciprocal inhibition of MeCP2 and LEDGF dimer formation. Following AlphaScreen titrations were performed: **A.** Increasing concentrations of Flag-MeCP2 E2 using Flag-donor and Flag-acceptor beads. **B**. Fixed concentration of 5 nM Flag-MeCP2 E2 and outcompetition with increasing concentrations of His6-ID-TRD using Flag-donor and Flag-acceptor beads. **C**. Fixed concentration of 5 nM Flag-MeCP2 E2 and outcompetition with increasing concentrations of His6-ID-TRD WT or R306C mutant using Flag-donor and Flag-acceptor beads. **D**. Fixed concentration of 5 nM Flag-MeCP2 E2 and outcompetition with increasing concentrations of GST-LEDGF/p52 using Flag-donor and Flag-acceptor beads. **E**. Representative AlphaScreen cross titration between GST-LEDGF/p52 and Flag-LEDGF/p75. **F.** Fixed concentration of 50 nM GST-LEDGF/p52 + 50 nM Flag-LEDGF/p75 and increasing concentrations of His6-MeCP2 E2. Error bars represent the SD; n = 3. A non-linear regression – sigmoidal curve fit with 1/Y^2^ weighting was fitted to the AlphaScreen data. See also Figure S1.

Previous studies showed that LEDGF also forms dimers based on electrostatic interactions^30,31^. We performed an AlphaScreen cross titration with GST-LEDGF/p52 and Flag-LEDGF/p75, revealing the known interaction between both LEDGF isoforms (Figure 6E)^30,31^. A fixed concentration of 50 nM GST-LEDGF/p52 and 50 nM Flag-LEDGF/p75 was used to form LEDGF heterodimers. These were outcompeted by using increasing concentrations of His_6_-MeCP2 E2 (Figure 6F). In conclusion, these results indicate that MeCP2 multimers can be disrupted by LEDGF and vice versa.

### LEDGF modulates MeCP2 chromatin condensation

MeCP2 is known to form pericentromeric condensates inside the nucleus of NIH3T3 cells as a result of liquid-liquid phase separation^14^. These speckles are functionally important for the MeCP2-chromatin interaction and the condensation of heterochromatin. Endogenous mouse MeCP2 was stably depleted in NIH3T3 cells with a shRNA and rescued with human MeCP2-eGFP E1 or E2. The human MeCP2-eGFP displayed the typical speckled pattern as well in NIH3T3 cells (Figure 7A). Subsequently, both or individual isoforms of LEDGF were depleted in the MeCP2-eGFP E1 or E2 NIH3T3 cells (Figure 7B). In order to assess the effect of LEDGF depletion on MeCP2 speckle formation and chromatin condensation, the cells were synchronized using 10 µg/mL aphidicolin and MeCP2 and heterochromatin speckles were imaged at different time points using the Operetta CLS High Content imager.

**Figure 7:**
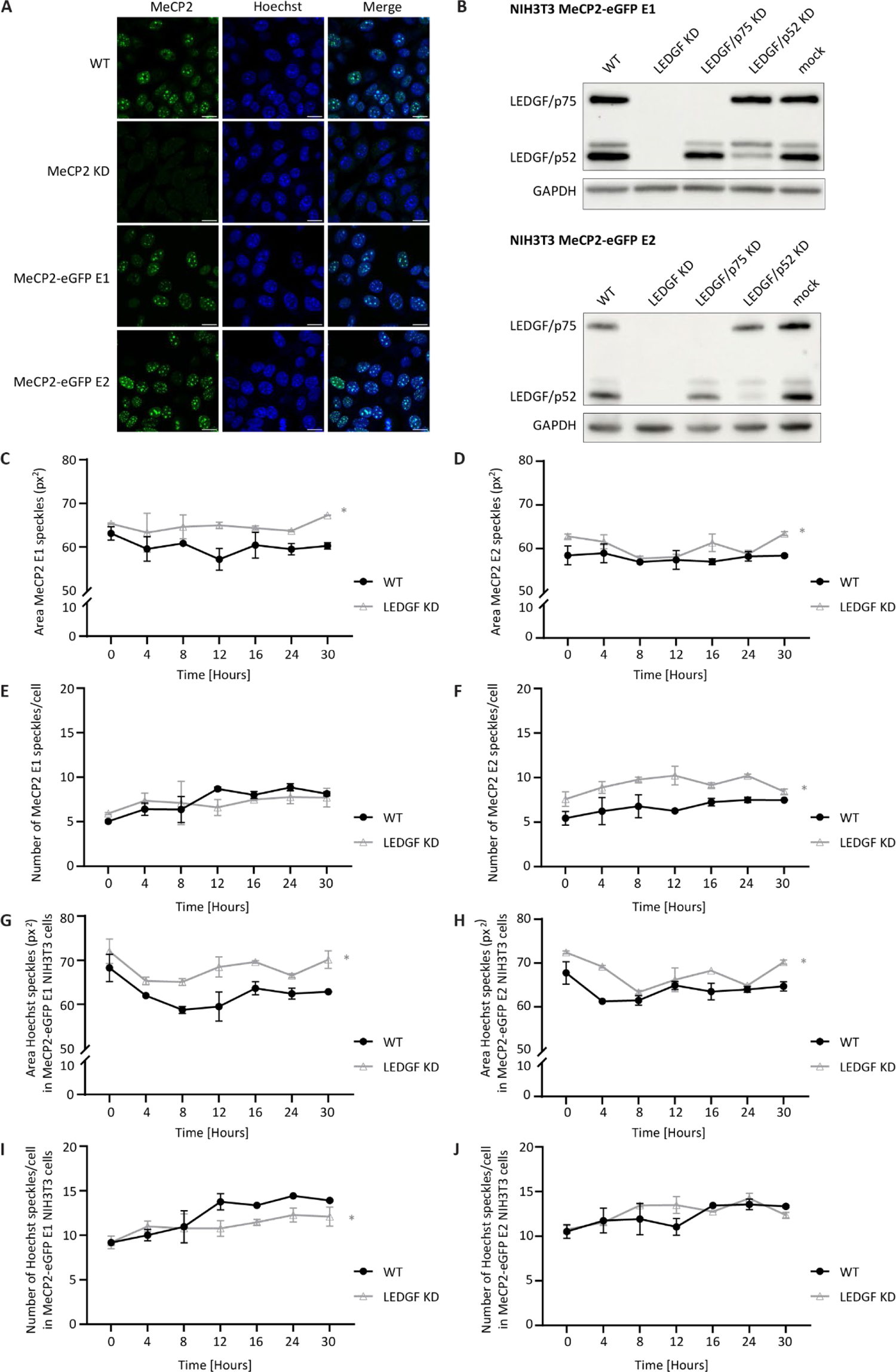
LEDGF depletion enlarges MeCP2 condensates and increases heterochromatin. **A.** Confocal microscopy images show distribution of MeCP2 in NIH3T3 cells. Endogenous mouse MeCP2 was detected in WT and MeCP2 KD NIH3T3 cells using a rabbit anti-MeCP2 antibody (1:500) and an anti-rabbit Alexa fluor 488 secondary antibody (1:1000). Human MeCP2-eGFP was detected in the 488 channel (green) and Hoechst in the 405 channel (blue). Scale bar is 20 µm. **B.** Western blot of LEDGF KD in MeCP2-EGFP E1 and E2 in NIH3T3 cells. LEDGF was detected with an anti-LEDGF-PWWP antibody (1:1000). GAPDH was detected with an anti-GAPDH-antibody (1:1000). **C**. Mean area of MeCP2-eGFP E1 speckles. **D**. Mean area of MeCP2-eGFP E2 speckles. **E.** Mean number of MeCP2-eGFP E1 speckles/cell. **F.** Mean number of MeCP2-eGFP E2 speckles/cell. **G.** Mean area of Hoechst speckles in MeCP2-eGFP E1 NIH3T3 cells. **H.** Mean area of Hoechst speckles in MeCP2-eGFP E2 NIH3T3 cells. **I.** Mean number of Hoechst speckles/cell in MeCP2-eGFP E1 NIH3T3 cells. **J.** Mean number of Hoechst speckles/cell in MeCP2-eGFP E2 NIH3T3 cells. Samples of NIH3T3 WT and LEDGF KD cells were measured over time. All measurements were performed using the Operetta CLS High Content Analysis System. Mean area is represented as number of pixels. Data of a representative experiment is shown as mean ± SD; n = 2. Statistical analysis was done using two-way ANOVA followed by Dunnett’s multiple comparison test vs. WT (* p < 0.05). See also Figure S2.

The area of MeCP2-eGFP speckles was significantly larger when both LEDGF/p75 and LEDGF/p52 were depleted in the cells (Figure 7C-D, Table 1). This increase was observed for both MeCP2-eGFP E1 (Figure 7C) and for MeCP2-eGFP E2 (Figure 7D) with 5% and 8% larger condensates compared to WT, respectively. A 37% increase in the number of MeCP2-eGFP speckles was seen for MeCP2 E2 (Figure 7F), although this phenotype was not observed for MeCP2-eGFP E1 (Figure 7E, Table 1). We also observed an impact on the area and size of MeCP2-eGFP speckles when either LEDGF/p75 or LEDGF/p52 was depleted, but to a lesser extent (Figure S2A-D). Taken together, depletion of LEDGF led to more crowding of MeCP2, resulting in larger and/or more MeCP2 condensates in the cell.

**Table 1:** Effect of LEDGF depletion on MeCP2 condensates.

|  | Number of MeCP2 speckles<sup>a</sup> | Area of MeCP2 speckles<sup>b</sup> | Number of Hoechst speckles<sup>a</sup> | Area of Hoechst speckles<sup>b</sup> |
| --- | --- | --- | --- | --- |
| **MeCP2-eGFP E1 NIH3T3** |  |  |  |  |
| WT | 7.36 ± 1.46 | 60.1 ± 2.25 | 12.23 ± 2.13 | 62.46 ± 3.30 |
| LEDGF/p75 KD | 8.39 ± 1.44 * | 59.95 ± 2.09 | 12.53 ± 1.96 | 64.02 ± 2.56 * |
| LEDGF/p52 KD | 7.72 ± 0.92 | 62.89 ± 1.72 * | 12.09 ± 1.29 | 62.98 ± 1.90 |
| LEDGF KD | 7.14 ± 1.05 | 64.77 ± 1.92 * | 11.08 ± 1.20 * | 68.14 ± 2.80 * |
| **MeCP2-eGFP E2 NIH3T3** |  |  |  |  |
| WT | 6.7 ± 0.96 | 57.88 ± 1.33 | 12.23 ± 1.39 | 63.92 ± 2.37 |
| LEDGF/p75 KD | 7.07 ± 0.97 | 60.37 ± 1.97 * | 11.58 ± 1.43 * | 66.88 ± 2.84 * |
| LEDGF/p52 KD | 7.41 ± .95 * | 60.46 ± 1.45 * | 12.12 ± 1.14 | 66.6 ± 2.50 * |
| LEDGF KD | 9.18 ± 1.04 * | 60.51 ± 2.37 * | 12.66 ± 1.21 | 67.76 ± 3.16 * |

Cells were synchronized using 10 µg/mL aphidicolin. MeCP2-eGFP E1 or E2 and heterochromatin speckles were imaged using the Operetta CLS High Content imager. Samples were measured at 0h, 4h, 8h, 12h, 16h, 24h and 30h. Data of a representative experiment is shown as mean ± SD; n = 2. Statistical analysis was done using two-way ANOVA followed by Dunnett's multiple comparison test vs. WT (\* p < 0.05). <sup>a</sup> Mean number of Hoechst or MeCP2-eGFP speckles per nucleus over time. <sup>b</sup> Mean area of Hoechst or MeCP2-eGFP speckles per nucleus over time represented as number of pixels.

As MeCP2 speckles in NIH3T3 cells colocalize with heterochromatin, visualized with Hoechst staining (Figure 7A), we next assessed the effect of LEDGF depletion on heterochromatin speckles. LEDGF KD resulted in significantly larger heterochromatin condensates (Figure 7G-H, Table 1). For MeCP2-eGFP E1 the heterochromatin area increased with 9% and for MeCP2-eGFP E2 with 8%. An effect of LEDGF depletion on the number of Hoechst speckles was not clearly observed as only a small reduction was observed in MeCP2-eGFP E1 NIH3T3 cells (Figure 7I-J, Table 1). In analogy to the observations with MeCP2-eGFP speckles, the effect of LEDGF depletion on the size and area of Hoechst speckles was less pronounced when only one LEDGF isoform was depleted (Figure S2G-H). In summary, a strong correlation on the effect of LEDGF depletion on either MeCP2-eGFP or Hoechst speckles was evidenced. LEDGF depletion increased both MeCP2 condensation and heterochromatin formation.

## Discussion

The findings presented in this study contribute to a deeper understanding of the molecular interaction between MeCP2 and LEDGF, shedding light on the biological relevance. The interaction between MeCP2 and LEDGF had been explored in previous studies, but the domains responsible for this interaction remained unknown. This study aimed to elucidate the specific interaction domains of MeCP2 and LEDGF, as well as to investigate the functional implications of MeCP2-LEDGF complex formation in the cell.

### Characterization of the MeCP2-LEDGF interaction

The interaction between MeCP2 and LEDGF was confirmed through a series of co-IP and AlphaScreen experiments with recombinant proteins (Figure 2). Our results corroborate the earlier studies on the interaction between MeCP2 and LEDGF^16,17^. By using deletion constructs of LEDGF, we confirmed that the interacting domain in LEDGF is located in the N-terminal region that is shared between the two isoforms (Figure 2F, Figure 4I, Figure S1C). More specifically, the PWWP-CR1 region of LEDGF is responsible for the interaction with MeCP2 as it was earlier suggested by Leoh *et al.*^17^. One key finding in our study is that both isoforms of LEDGF interacted with both isoforms of MeCP2, yet LEDGF/p52 showed an apparent higher affinity for MeCP2 than LEDGF/p75 in co-immunoprecipitation experiments (co-IP; Figure 2A-B). In MeCP2, on the other hand, the ID-TRD domain proved important for the interaction with LEDGF (Figure 3-4). This result was expected as the ID-TRD of MeCP2 is a known interaction site for many MeCP2 interaction partners^1^.

MeCP2 is an intrinsically disordered protein^32,33^. Characteristically, these proteins lack a stable three-dimensional structure and interact electrostatically with DNA and other proteins. In MeCP2 only the MBD is structured, while the ID and TRD are structurally disordered but form a secondary structure upon binding interaction partners and/or DNA^1,34^. We postulated an electrostatic interaction between the ID-TRD and LEDGF which is often the case for structurally disordered proteins^32,33^. Three positively charged regions are present in the TRD and we mutated the four or five positively charged residues to alanines (Figure 5A). Using this approach, we could narrow down the interaction site in MeCP2 to the C-terminal residues of the TRD. More specifically, the positively charged amino acids 266-309 appeared to be important for the interaction (Figure 5C-D). This result is also consistent with the PWWP-CR1 interaction domain in LEDGF wherein the CR1 domain is unstructured and carries regions of negatively charged residues.

The C-terminal region of the TRD, also known as the NID, is the interaction site of another important MeCP2 binding partner: the NCoR complex^4,35^. The positively charged cluster 302-306 of conserved amino acids represents a recruitment surface for the NCoR complex^4^. R306C is a known clinical Rett syndrome (RTT) mutant and is the most common missense mutation in the NID of MeCP2 (Figure 5A-B)^27^. We observed a clear reduction in the binding affinity of the R306C MeCP2 mutant for LEDGF (Figure 5E). Although R306C is also defective for interaction with NCoR, a role for LEDGF in the pathophysiology of RTT patients with the R306C mutation may be plausible. Targeting the MeCP2-LEDGF interaction may prove a novel therapeutic strategy.

DNA Methyltransferase 3A (DNMT3A) is also known to interact with MeCP2 through the TRD^36^. While DNMT3A is known to directly bind MeCP2 residues 214-228, mutations in this region of MeCP2 only showed a minimal reduction in LEDGF binding, indicating that DNMT3A and LEDGF do not compete for the same residues. However, it remains possible that a competition exists due to steric hindrance or that these residues support the proper folding of MeCP2 upon interaction with LEDGF. Interestingly, while DMNT3A also contains a PWWP domain, the chromatin interaction ADD domain of DNMT3a and not the PWWP domain is responsible for the interaction with MeCP2^36^.

The interaction between MeCP2 and LEDGF showed to be strongly DNA-dependent (Figure 2E). Since both are DNA-binding proteins, indirect complex formation may occur due to DNA-bridging. Additionally, the presence of DNA may also affect the protein folding of unstructured domains in MeCP2 and LEDGF. As the ID-TRD of MeCP2 is prone to non-specific DNA interactions^34^, we had to exclude that the interaction with LEDGF occurs also in the absence of DNA (Figure 4C,H). We observed a residual interaction after MNase treatment, indicating a direct MeCP2-LEDGF interaction independent from DNA.

### Impact of LEDGF on MeCP2 function

The disordered structure of proteins like MeCP2 supports their functional versatility. Interactions are context-dependent which enables MeCP2 to participate in various cellular processes. MeCP2 is known to form electrostatic self-interactions through the ID-TRD domain^28^. These self-interactions were shown to occur in the absence of DNA^28^. We confirmed that MeCP2 forms electrostatic self-interactions through its ID-TRD domain (Figure 6A-B, Figure S1F). Interestingly, the R306C mutation did not affect MeCP2 multimerization (Figure 6C). This result suggests that MeCP2 and LEDGF do not compete for exactly the same amino acid residues in the ID-TRD domain of MeCP2. However, steric hindrance may cause competition for binding.

MeCP2 is a very versatile protein. One of the most described functions of MeCP2 is transcriptional repression and activation. However, the molecular mechanism by which MeCP2 regulates gene expression remains poorly understood^37,38^. An increasing number of studies reveal that transcriptional profiling of RNA in mice lacking functional MeCP2 do not reveal significant gene expression changes^39–42^. These studies show evidence for large numbers of subtle alterations, comprising both increases and decreases in gene expression. The majority of changes had a magnitude less than 50%, thus making it difficult to distinguish between biological changes from background^43^. The genome-wide distribution of MeCP2 is one of the important aspects to consider in these observations. When a transcriptional regulator affects a small number of genes, significant expression changes are expected when the transcriptional regulator is deficient. However, if a transcriptional regulator regulates a large number of genes, it is more difficult to predict how deficiency will impact gene expression as the supply of transcriptional machinery is limited in the cell. It has been hypothesized that the MeCP2-LEDGF interaction may play a role in modulating transcriptional activity^17^. Considering that MeCP2 is globally distributed in the nucleus and that LEDGF interacts with the RNA polymerase II complex^44^, it is possible that the MeCP2-LEDGF interaction is involved in transcriptional regulation. A MeCP2-LEDGF complex might have implications for the activation or repression of a wide range of MeCP2 regulated genes. Since in previous studies RNA-seq experiments on MeCP2 deficient models did not reflect these broad effects of MeCP2 on transcription, future studies might require more sensitive methods to detect gene expression alterations.

It has been shown that changes in MeCP2 expression cause large-scale chromatin reorganization^45^. MeCP2 homo-interactions appear to be essential for heterochromatin organization^15,28,46–49^. Results from our MeCP2 speckle assay show that LEDGF depletion increases MeCP2 condensation in the cells (Figure 7, Figure S2). Additionally, we show that LEDGF can disrupt MeCP2 dimers (Figure 6D) and that the absence of LEDGF increases MeCP2 condensation and heterochromatin formation (Figure 7C-F). Our results show that LEDGF may play a role in heterochromatin formation by modulating MeCP2 oligomerization. Previously, a model has been proposed in which a single MeCP2 simultaneously binds two nucleosomes to form a ‘sandwich’ that forms chromatin loops and compacts DNA^15^. We propose instead that MeCP2 forms a dimer or multimer through its ID-TRD domain, allowing each MBD domain to bind nucleosomes in order to form chromatin loops and compact DNA. Methods to determine genome-wide chromatin status, such as ATAC-seq, will provide valuable information to support the results of this study.

### Impact of MeCP2 on LEDGF function

While LEDGF modulates MeCP2 condensates, MeCP2 by itself can disrupt LEDGF dimers (Figure 6F). Even though the function of LEDGF dimers is not fully understood, disruption of LEDGF dimers may impact the various functions of LEDGF in the cell, such as transcriptional regulation and DNA damage repair^18,21,44,50–52^. Considering that both MeCP2 and LEDGF are known to be involved in transcriptional regulation, MeCP2 may conversely affect LEDGF-dependent gene regulation. It has been previously hypothesized that MeCP2 dampens transcriptional noise throughout the genome^53^. This is supported by the fact that MeCP2 deficiency leads to higher transcriptional noise from repetitive elements, such as LINE-1 retrotransposons^53–55^. The binding of MeCP2 to LEDGF may have a dampening effect on LEDGF-dependent gene regulation, thereby affecting transcriptional activation by LEDGF. Trans-activation assays will have to show whether MeCP2 can influence LEDGF-dependent gene regulation to support this hypothesis.

As both isoforms of LEDGF are capable of binding MeCP2 (Figure 2A-C), the functional difference between LEDGF/p75 and LEDGF/p52 may also affect their respective function when bound to MeCP2. LEDGF/p75 is known to tether via its IBD-domain, which is not present in LEDGF/p52, several proteins to the chromatin. LEDGF/p75 is known to bind transcriptional regulators such as JPO2, PogZ, MED1, IWS1 and MLL and is also involved in DNA-damage repair by recruiting proteins from the homologous recombination repair pathway^22,52,56,57^. In contrast, not much is known about the specific function of LEDGF/p52. One study suggested a specific role for LEDGF/p52, and not LEDGF/p75, in modulating splicing due to its colocalization and interaction with mRNA processing proteins^58^. The differential roles of LEDGF/p52 and LEDGF/p75 raise the possibility that the biological role of the MeCP2-LEDGF/p52 complex differs from that of the MeCP2-LEDGF/p75 complex.

LEDGF is known to bind di- and trimethylated H3K36 marks, while MeCP2 is known to bind methylated histone H3K9 and H3K27 marks. A study by Lee *et al.* showed a correlation between the impact of MeCP2 on transcription and binding of MeCP2 to H3K27 histone marks^13^. Regardless of the fact that they do not bind the same histone modifications, steric hindrance for nucleosome binding may cause competition between MeCP2 and LEDGF. While competition for the nucleosomes would not necessarily involve a direct MeCP2-LEDGF interaction, it could have functional implications on transcriptional regulation.

MeCP2 is highly expressed in the brain and is essential for the development and maintenance of neurons^59^. Previous studies have shown that MeCP2 expression levels are tightly controlled to maintain proper neuronal function and overall cellular homeostasis. The occurrence of RTT and MeCP2 Duplication Syndrome (MDS), two neurodevelopmental disorders which are characterized by too little or too much functional MeCP2, respectively, also support this idea. The results of our study suggest that MeCP2 and LEDGF may contribute to a finely tuned regulatory system in the cell in which the balance between MeCP2 and LEDGF is key. Our study suggest that further research on the functional role of the MeCP2-LEDGF interaction is needed, potentially offering new avenues for therapeutic interventions in RTT and MDS.

## Acknowledgements

Financial support was received from CELSA (KU Leuven) CELSA5439-DOA/18/003 and Fonds Wetenschappelijk Onderzoek (EFH-D7808-G0C2620N FWO). Y.H. received a personal fellowship from Fonds Wetenschappelijk Onderzoek (1185621N). We thank the Cell and Tissue Imaging Core (KU Leuven) and the VIB Bioimaging Core (VIB-KU Leuven) for the use of their core facilities. All MeCP2 sequences were cloned from the pcDNA-Flag-MeCP2 plasmid, a kind gift from Prof. Dr. Adrian Bird (University of Edinburgh, Edinburgh, UK).

## Author Contributions

Z.D., S.L., S.V.B., R.L., and Y.H. conceptualized and designed experiments. S.L. wrote the original manuscript. S.L., S.V.B., R.L. and K.H. performed experiments and analyzed the data. P.V. and A.S. performed experiments and provided technical support. Z.D. and F.C. supervised the project and acquired funding. All authors have read and agreed to the published version of the manuscript.

## Data and Material Availability Statement

GraphPad Prism version 10.0 for Windows [GraphPad Software, San Diego, CA, USA] was used for data analysis and graph representation. Structures were made in PyMol Molecular Graphics System [Schrödinger, LLC]. Figures were made in Adobe Illustrator. Additional data and material generated in this study are available from the corresponding author Zeger Debyser.

