## Supplementary Files for "LEDGF interacts with the NID of MeCP2 and modulates MeCP2 condensates"

### Supplementary Figures

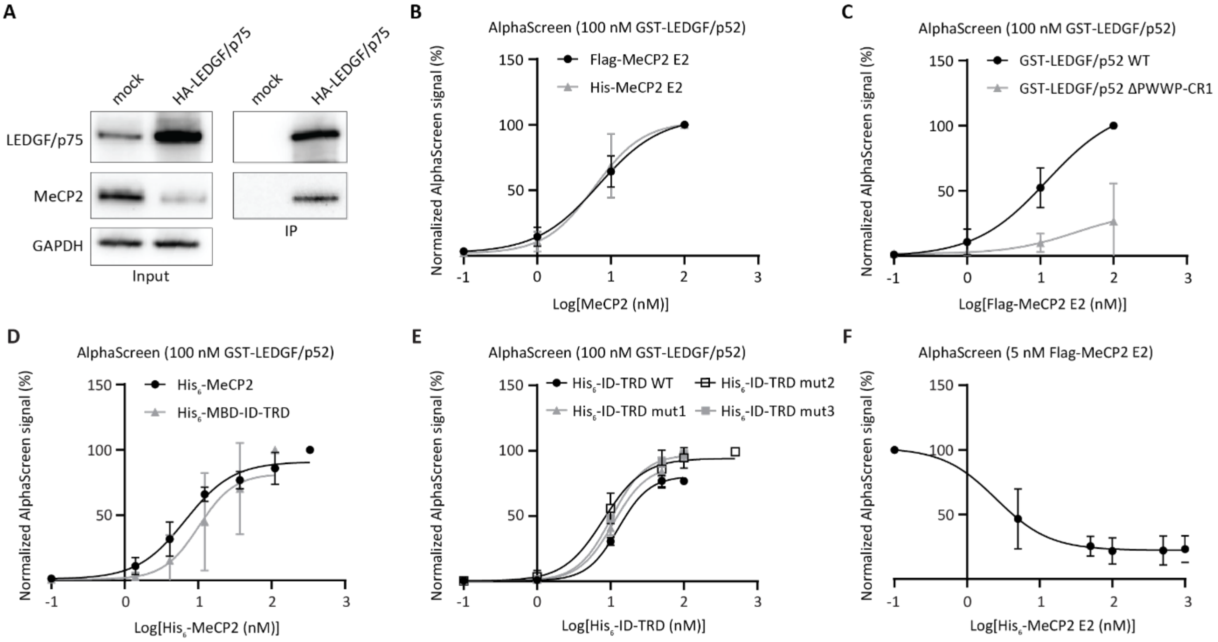


***Figure S1: MeCP2 and LEDGF co-immunoprecipitate and interact in AlphaScreen assays. A****. Co-IP of LEDGF/p75. HEK293T cells were transfected with a plasmid encoding HA-LEDGF/p75. 24 hours after transfection anti-HA-beads were used for immunoprecipitation of the cell lysate. Precipitated proteins were analyzed on western blot. LEDGF was detected with an anti-LEDGF-PWWP antibody (1:1000). MeCP2 was detected with an anti-MeCP2 antibody (1:500). GAPDH was detected with an anti-GAPDH-antibody (1:1000). Following AlphaScreen titrations were performed:* ***B.*** *100 nM GST-LEDGF/p52 and increasing concentrations of Flag-MeCP2 E2 or His-MeCP2 E2. AlphaScreen counts were first normalized within each condition to avoid confounding for the use of different AlphaScreen beads.* ***C.*** *100 nM GST-LEDGF/p52 WT or ΔPWWP-CR1 and Flag-MeCP2 E2.* ***D.*** *100 nM GST-LEDGF/p52 and increasing concentrations of His-MeCP2 E2 or His-MBD-ID-TRD without MNase treatment.* ***E.*** *100 nM GST-LEDGF/p52 and increasing concentrations of His-ID-TRD WT or mutants without MNase treatment.* ***F.*** *Fixed concentration of 5 nM Flag-MeCP2 E2 and outcompetition with increasing concentrations of His-MeCP2 E2. Error bars represent the SD; n = 3. A non-linear regression – sigmoidal curve fit with 1/Y² weighting was fitted to the data.*

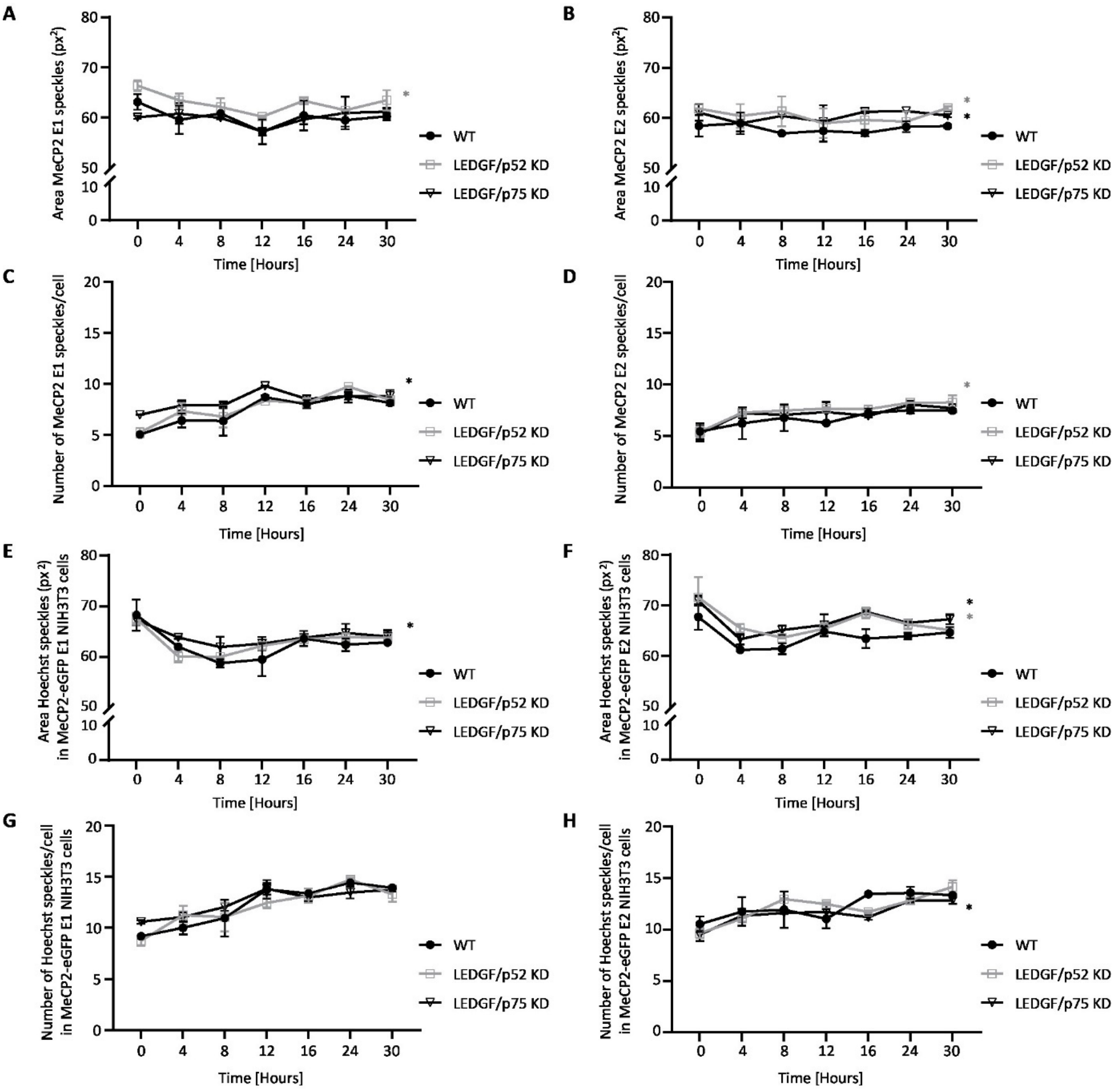


***Figure S2: LEDGF depletion enlarges MeCP2 condensates and increases heterochromatin. A.*** *Mean area of MeCP2-eGFP E1 speckles.* ***B****. Mean area of MeCP2-eGFP E2 speckles.* ***C.*** *Mean number of MeCP2-eGFP E1 speckles/cell.* ***D.*** *Mean number of MeCP2-eGFP E2 speckles/cell.* ***E.*** *Mean area of Hoechst speckles in MeCP2-eGFP E1 NIH3T3 cells.* ***F.*** *Mean area of Hoechst speckles in MeCP2-eGFP E2 NIH3T3 cells.* ***G.*** *Mean number of Hoechst speckles/cell in MeCP2-eGFP E1 NIH3T3 cells.* ***H.*** *Mean number of Hoechst speckles/cell in MeCP2-eGFP E2 NIH3T3 cells. Samples of NIH3T3 WT, LEDGF/p52 and LEDGF/p75 KD cells were measured over time. All measurements were performed using the Operetta CLS High Content Analysis System. Mean area is represented as number of pixels. Data of a representative experiment is shown as mean ± SD; n = 2. Statistical analysis was done using two-way ANOVA followed by Dunnett's multiple comparison test vs. WT (* p < 0.05).*

***Table S1: Target sequences for MeCP2 and LEDGF shRNAs.***

| shRNA target | Sequence |
|---|---|
| MeCP2 E1 and MeCP2 E2 | tgacaaagcttcccgattaac |
| LEDGF/p75 and LEDGF/p52 | aagcaatgaggatgtgactaaa |
| LEDGF/p75 | agacagcatgaggaagcgaatta |
| LEDGF/p52 | aagcaccaaacaacatgtaatc |
